# A latitudinal pattern of plant leaf-associated bacterial community assembly

**DOI:** 10.1101/2022.06.23.497395

**Authors:** Zihui Wang, Yuan Jiang, Minhua Zhang, Chengjin Chu, Yongfa Chen, Shuai Fang, Guangze Jin, Mingxi Jiang, Juyu Lian, Yanpeng Li, Yu Liu, Keping Ma, Xiangcheng Mi, Xiujuan Qiao, Xihua Wang, Xugao Wang, Han Xu, Wanhui Ye, Li Zhu, Yan Zhu, Fangliang He, Steven W. Kembel

**Affiliations:** Département des sciences biologiques, Université du Québec à Montréal, Montréal, Québec, H2X 1Y4, Canada; ECNU-Alberta Joint Lab for Biodiversity Study, Tiantong Forest Ecosystem National Observation and Research Station, School of Ecology and Environmental Sciences, East China Normal University, Shanghai, 200241, China; Department of Renewable Resources, University of Alberta, Edmonton, Alberta, T6G 2H1, Canada; State Key Laboratory of Biocontrol, School of Life Sciences / School of Ecology, Sun Yat-sen University, Guangzhou, 510275, China; Center for Ecological Research, Northeast Forestry University, Harbin, 150040, China; Key Laboratory of Aquatic Botany and Watershed Ecology, Wuhan Botanical Garden, Chinese Academy of Sciences, Wuhan, 430074, China; Key Laboratory of Vegetation Restoration and Management of Degraded Ecosystems, South China Botanical Garden, Chinese Academy of Sciences, Guangzhou, 510650, China; State Key Laboratory of Vegetation and Environmental Change, Institute of Botany, Chinese Academy of Sciences, Beijing, 100093, China; Zhejiang Tiantong Forest Ecosystem National Observation and Research Station, School of Ecology and Environmental Sciences, East China Normal University, Shanghai, 200241, China; CAS Key Laboratory of Forest Ecology and Management, Institute of Applied Ecology, Chinese Academy of Sciences, Shenyang, 110016, China; Research Institute of Tropical Forestry, Chinese Academy of Forestry, Guangzhou, 510520, China

## Abstract

Plant-associated microbes are essential for promoting plant well-being, maintaining biodiversity, and supporting ecosystem function. However, little is known about the geographic distribution of plant-microbe symbioses and how they are formed and change along latitudinal gradients. Here we identified leaf bacteria for 328 plant species sampled from 10 forests along a tropical to temperate gradient in China. We analyzed the diversity and composition of plant leaf-associated bacteria and quantified the contributions of hosts, habitats, and neighborhood plants to the plant-bacterial symbiosis. We found a strong latitudinal gradient in leaf bacterial diversity and composition. Bacterial assemblages on leaves were most strongly selected by host plants, and the selection pressure increased with latitude. In contrast, at low latitudes and at large geographical scales multiple factors were found to jointly regulate bacterial community composition. Our result also showed that plant-bacteria symbiotic networks were structured by network hub bacteria taxa with high co-occurrence network centrality, and the abundance of temperate hub taxa was more influenced by host plants than that in tropical forests. For the first time, we documented a previously unrecognized latitudinal gradient in plant-bacterial symbioses that was regulated by a joint effect of multiple factors at low latitudes but mostly by host selection at high latitudes, implying that leaf microbiomes are likely to respond differently to global change along the latitudinal gradient.

## Main text

All animals and plants are colonized by microorganisms, which are critical for ecosystem function and global biogeochemical cycles^1^. As one of the most common types of biotic interactions, plant-microbial symbioses are essential for plant growth, health and stress resilience, and also for supporting ecosystem functions including enhancing productivity, biogeochemical cycling, and diversity maintenance^2,3^. However, our knowledge about plant-microbial symbioses is highly variable among symbiosis types, with belowground symbioses such as nitrogen-fixation symbioses and root mycorrhizal symbioses being extremely well studied while aboveground plant-microbe symbioses have not been explored as deeply^4,5^. An example of aboveground symbioses is microbial assemblages on leaves – a metasymbiosis that is formed not by individual microbes or a specific group of microbes (i.e. *Bradyrhizobium* bacteria or glomeromycete mycorrhizal fungi in the case of roots) but by the diverse metacommunity of microorganisms available to colonize leaves^6^. How such leaf-microbe associations are formed and maintained is a critical but unanswered question, leading to a major concern that the future role of plant microbiomes in supporting plant well-being and sustaining ecosystem functioning may be compromised in the face of global change^7,8^. To address this question, it is necessary to understand the geographic variation of the phyllosphere metasymbiosis across a broad geographic scale in order to gain insights into how leaf-microbe associations are assembled and how they vary across different habitat conditions and host compositions. This understanding of geographic patterns of phyllosphere microbial diversity is indispensable for predicting the responses of plant microbiomes to global change^7,8^.

Bacteria are the most abundant microbes in the phyllosphere with an estimated 10^7^ bacterial cells per cm^2^ of leaf surface^9^. Considering the total area of plant leaf surfaces on Earth, the global bacterial population in the phyllosphere is enormous (up to 10^26^ cells), as is their impact on plant communities and ecosystem functions through effects including altering plant immunity, modifying plant hormone production and facilitating global carbon and nitrogen cycling^10–12^. The assembly of this ubiquitous bacterial leaf symbiosis is governed by a range of factors that influence the community assembly processes such as selection and dispersal^13^. For example, host plant attributes including species identity, functional traits and phylogeny shape bacterial community assembly through selection processes^14–16^. Neighborhood plants also select for and create a bacterial metacommunity that alters the composition of local communities through dispersal^17,18^. Additionally, abiotic environments such as temperature and microhabitats which are often spatially autocorrelated have been observed to affect leaf bacterial diversity^19–21^. However, these drivers of leaf bacterial assembly have commonly been examined separately in previous studies. As a result, there is poor understanding of the relative importance of these different drivers of plant-bacterial symbioses. Furthermore, current knowledge on the drivers of plant-microbe associations is almost exclusively based on individual sites; we understand very little how assembly processes of bacteria community vary along latitudinal gradients, in stark contrast with our knowledge about latitudinal diversity gradients of higher plants and animals^22,23^.

Studies of microbial co-occurrence networks have revealed hub taxa that occupy central positions within microbial interaction networks, and have demonstrated that these hub taxa play key roles driving microbial community structure and function^24,25^. At the metacommunity level, hub taxa connect local communities via dispersal and gene flow and influence local community assembly by microbe-microbe and plant-microbe interactions, through which their population dynamic could impact large-scale patterns of plant-bacterial interactions^26–28^. Identifying the hub taxa in plant-bacterial association at the metacommunity level is a pressing need given their importance in mediating eco-evolutionary feedbacks, but has been limited to date by a lack of large-scale datasets on plant-microbe associations^29^. Furthermore, knowledge about the factors that control the geographic distribution of these hub bacterial taxa is essential for predicting future change in microbiome structure due to gain or loss of hub taxa, but is currently unavailable for plant-associated microbes^30^.

Here, we addressed these knowledge gaps by investigating geographic patterns of phyllosphere bacterial diversity, their network structure and assembly mechanisms along a large latitudinal gradient for a network of stem-mapped forest sites across China. We quantified bacterial communities using high-throughput sequencing of the 16S rRNA gene for a total of 1453 leaf samples from 328 plant species from 10 forest sites across major forest biomes in China. Specifically, we aimed to (1) document latitudinal patterns of leaf-associated bacterial diversity, (2) disentangle the effects of hosts, environments, neighborhood plants and spatial factors on the community assembly of phyllosphere bacteria along the latitudinal gradient and across geographic scales, and (3) construct plant-bacterial symbiosis networks and identify the metacommunity-level hub taxa that are instrumental to plant-bacteria associations; we further modeled the distribution of these bacterial hub taxa along the latitudinal gradient.

## Results

We identified 49254 16S rRNA gene amplicon sequence variants (ASV) from 1453 leaf samples, belonging to 237 bacterial orders and 386 families. *Proteobacteria* were the dominant phylum, accounting for 70% of the total abundance of sequences across all samples. *Beijerinckiaceae* (*Rhizobiales*) were most abundant in tropical and subtropical regions while *Sphingomonadaceae* (*Sphingomonadales*) were most abundant in temperate forests (Fig. S1).

ASV richness of phyllosphere bacteria decreased with latitude (Fig. 1a). There was strong evidence that ASV richness was correlated positively with the diversity of neighborhood plants, negatively with slope, and was linked with elevation and annual temperature in quadratic relationships (Fig. S2). We used global non-metric multidimensional scaling ordinations to visualize the variation in bacterial community composition and show clear clustering of bacterial communities by study sites and host taxonomy (Fig. 1b&S3). Permutational multivariate analysis of variance revealed strong evidence that climate, soil conditions, topography, neighborhood plant diversity and host traits all impact bacterial community assembly (Table S1). There was a higher similarity of phyllosphere bacterial communities on phylogenetically closely related plants and spatially closer samples, leading to host phylogenetic structure and spatial autocorrelation effects on bacterial community assembly at both local and large scales (Fig. S4).

**Figure 1.**
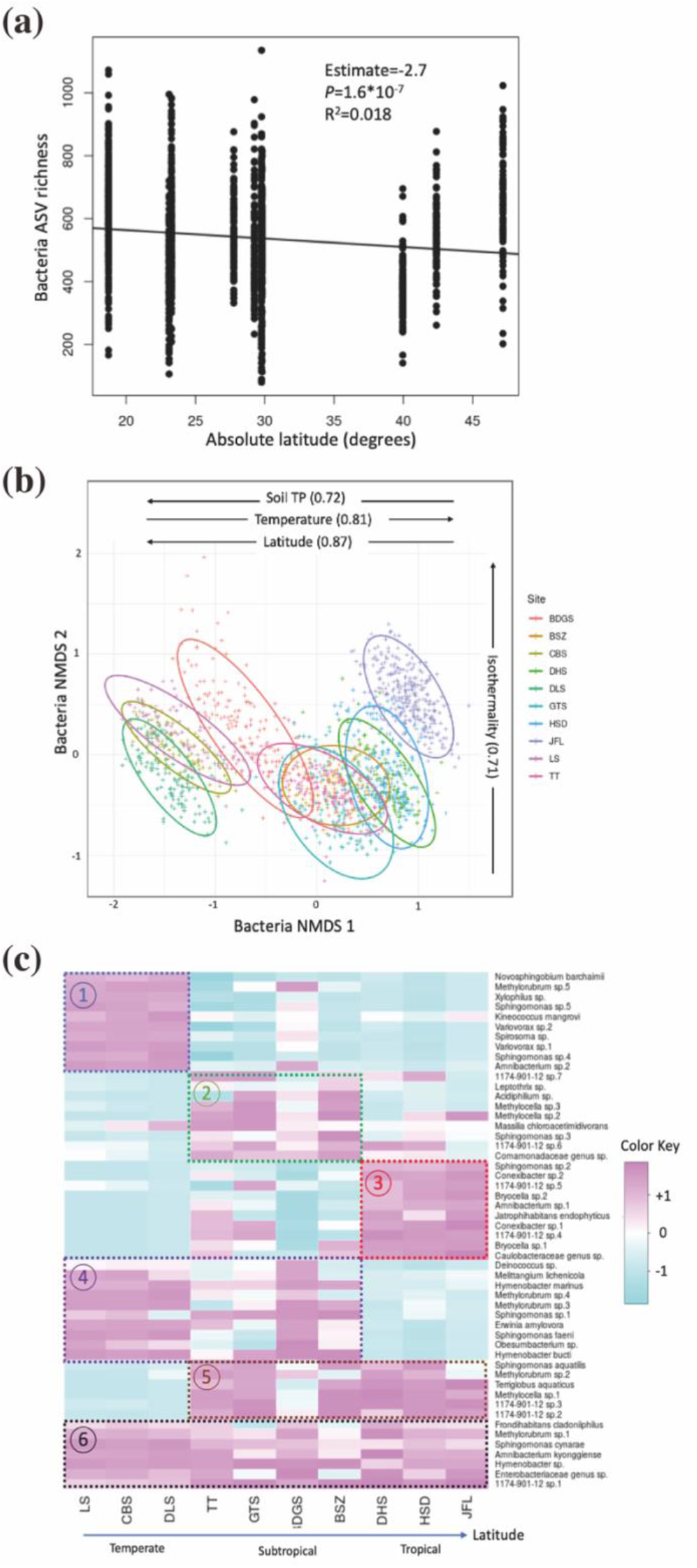
The latitude patterns of phyllosphere bacterial (a) diversity, (b) community composition and (c) occurrence of metacommunity-level hub ASVs. First-and second order polynomial fits are shown in black (dashed) and red (solid) respectively in (a), Bacterial community composition was visualized using nonmetric multidimensional scaling ordination with each ellipse representing site in (b), where arrows inside indicate significant variables correlated with ordination scores and the explained power (R^2^) of the correlation. Heatmap in (c) show the relative frequency of hub ASVs clustering regarding to (1) temperate hubs (2) subtropical hubs (3) tropical hubs, (4) hubs that connect temperate and subtropical, (5) hubs that connect subtropical and tropical sites, and (6) global hubs that connect all sites.

We quantified the variation in bacterial ASV richness and composition explained by multiple variables that have been proposed as drivers of host-associated microbial diversity using generalized linear models and redundancy analysis respectively. The variables included host-related factors (phylogenetic eigenvectors + plant traits), abiotic environment (soil conditions + topography + climate), diversity and abundance of neighborhood plants, and spatial eigenvectors (see Methods). At the local scale of individual sites, these models explained 26%-76% of the variation in ASV richness and 19%-36% of the variation in community composition of phyllosphere communities. At regional levels, these models explained 27%-68% of the variation in species richness, and 33%-42% of the variation in community composition respectively. At a ‘global’ scale (defined as the scale including all study sites), the model explained 43% of the total variation in phyllosphere bacterial species richness and 47% of the variation in community composition across all sites (Fig. 2&S5).

**Figure 2.**
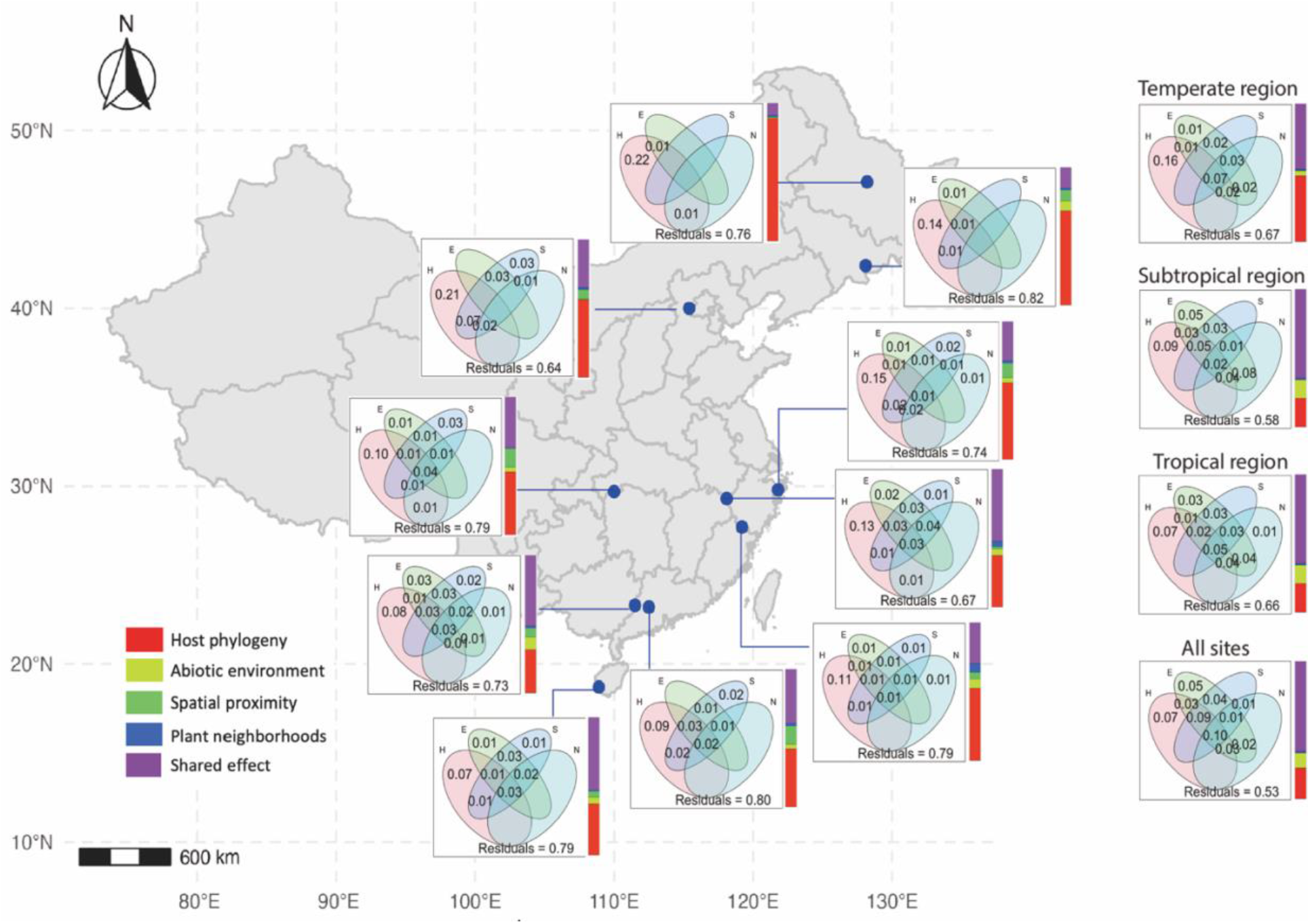
Map of study sites indicating the percentage of variation in community composition explained by host, abiotic environment, neighborhood plants and spatial factors at local scales for each site, regional scales for temperate, subtropical, and tropical sites, and a ‘global’ scale that includes all sites. The Venn diagrams show the absolute variance explained by each factor and the histograms on the right of Venn diagrams show the relative contribution of the independent and shared effects of four variables to the explained variance. Explained variation less than 1% was not shown in the Veen diagram. Abbreviation; H: host, E: abiotic environments; S: spatial variables; N: neighborhood plant communities.

We used variation partitioning to disentangle the independent and joint effect of these variables and quantify their relative importance in structuring bacterial diversity and community assembly. Generally, the independent effect of host was the most important in explaining both phyllosphere bacterial ASV richness and community composition, accounting for 15-89% of the explained variance, followed by abiotic environments and spatial eigenvectors (contributions of up to 29% and 25% respectively), while the attributes of plant neighborhoods were less important (only contributing 9% of explained variance). There was a strong joint effect of different variables, which accounted for 8-72% of the explained variation in bacterial community composition (Fig. 2). The relative importance of these factors varied across geographic scales and was highly latitude dependent; for example, the independent effect of host was positively correlated with latitude and was less important at large scale, while the joint effect of these four variables decreased with latitude and was more pronounced at large scales, and the overall contribution of host-related factors increased while that of other factors such as environment decreased with latitude (Fig. 3).

**Figure 3.**
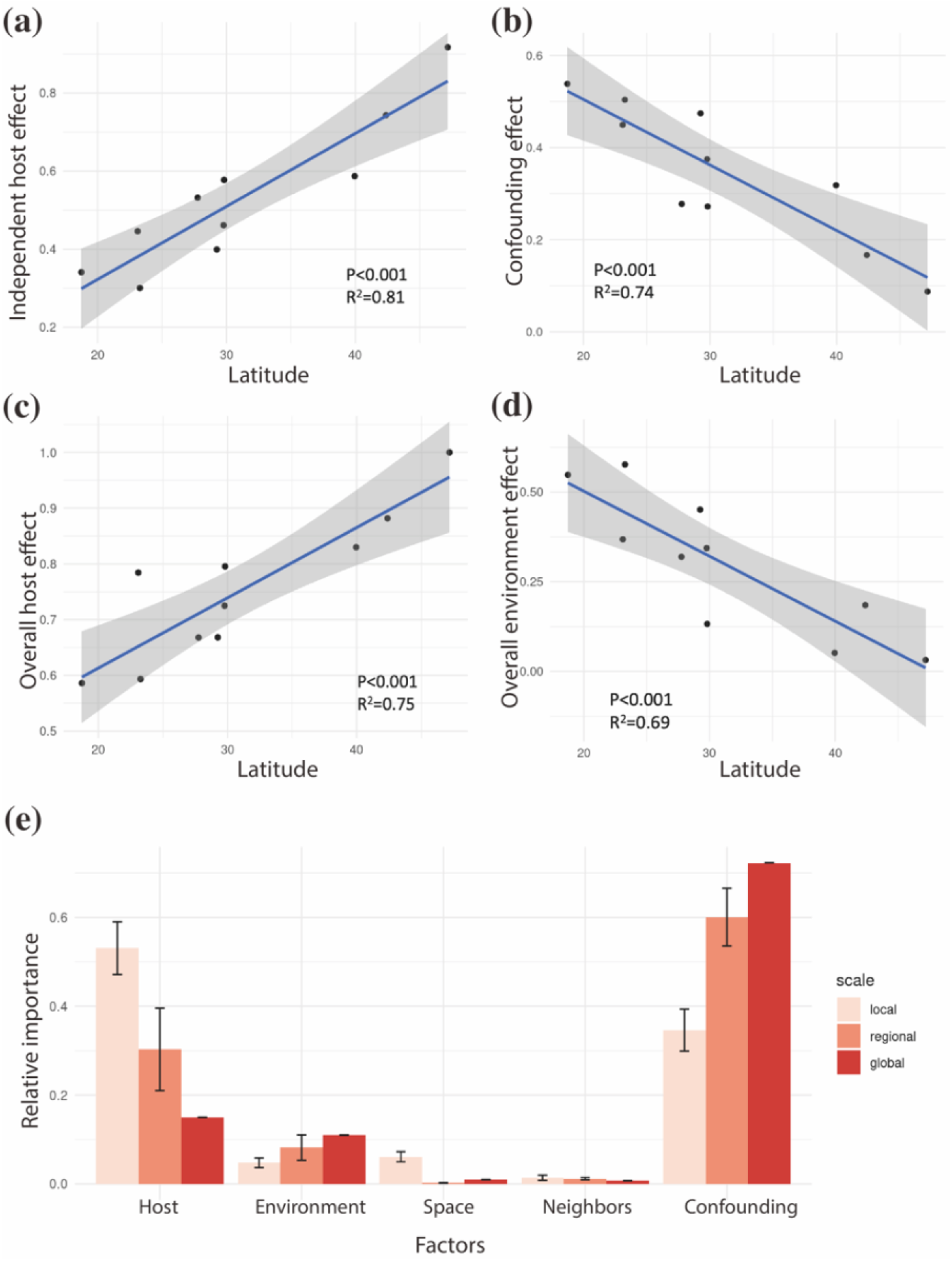
The relative importance of different mechanisms in structuring phyllosphere bacterial community assembly varies across latitude gradients (*a*-*d*) and across geographic scales (*e*). Coefficients of linear regression between the relative importance of drivers and latitude were reported in *a*-*d*, and Kruskal-Wallis rank sum test was applied to test the variation across three scales in *e*, showing moderate evidence (p=0.046) for increased confounding effects at larger scales (sample size: 10, 3 and 1 for local, regional and global scales).

We constructed metacommunity-level networks of plant-bacterial associations at regional and global scales and quantified the topological positions of ASVs within each co-occurrence network. Hub ASVs with high network betweenness centrality were identified for tropical, subtropical, and temperate regional networks and a global network respectively (see Methods). Hub bacterial ASVs differed across regions; in subtropical and tropical regions, bacteria hubs were primarily ASVs belonging to the *Rhizobiales* genera *1174-901-12, Methylocella* and *Methylobacterium*, while in temperate networks the hubs included *Sphingomonas* (*Sphingomonadales*) and *Hymenobacter* (*Cytophagales*) (Fig. S6). These bacterial ASVs connect with most of the nodes (i.e. plant species) in the association networks and structure plant-bacterial association networks at metacommunity levels; plant-phyllosphere bacteria association networks at local sites were interconnected by hub bacterial ASVs to form regional-scale networks, and these regional networks were further linked together through hub ASVs at global scales (Fig. 4).

**Figure 4.**
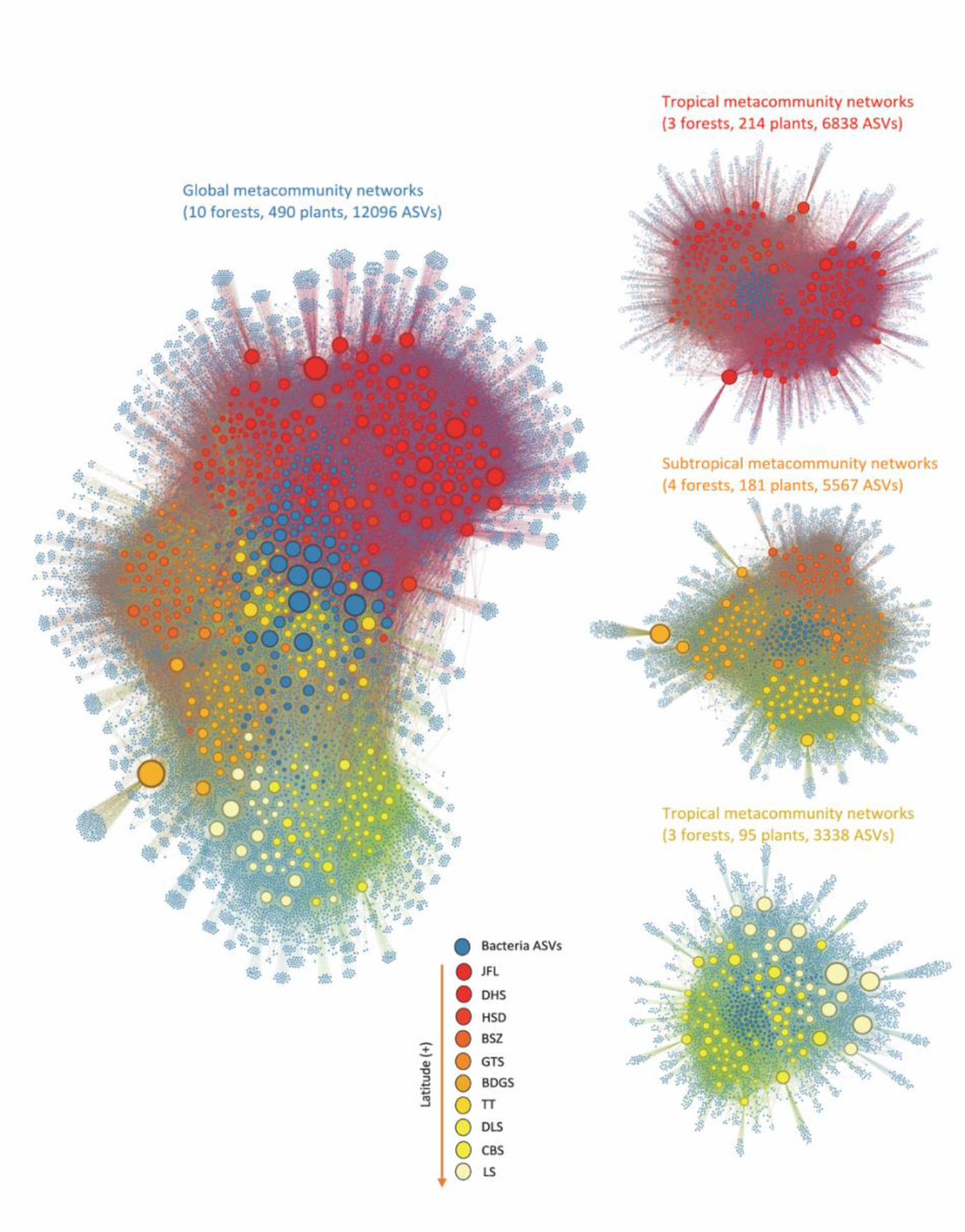
Global and regional networks representing the associations between plants and phyllosphere bacteria. Circles represent either bacteria ASVs (dark blue) or plant species (warm colors). A link was established if the plant species was associated with more than 15 reads of an ASV (see Methods). Size of the circles indicates the betweenness centrality of nodes. Plant species were colored according to latitude of site, including tropical: JFL, DHS, HSD; subtropical: BSZ, GTS, BDGS, TT; and temperate sites: DLS, CBS, LS.

The distribution of phyllosphere bacterial hubs was clustered along latitudinal gradients (Fig. 1c). To determine the relative importance of different factors structuring the biogeography of these hub ASVs, we applied a joint species distribution modeling approach where nine habitat variables were taken as fixed effects and two distance matrices representing the spatial and host phylogenetic structure as random effects (see Methods). The models explained up to 90% and 73% of the variation in presence-absence and abundance of hub bacterial ASV, respectively.

Overall, climate (including temperature, precipitation and isothermality) was the most important factor explaining the distribution of phyllosphere bacterial hub ASVs (Fig. 5&S7). There was also a major contribution of the random effects of spatial factors and host phylogeny on hub ASV abundance, highlighting the importance of taking spatial and host phylogenetic structure into account for unbiased inference and to improve the prediction of the distribution of plant microbiomes. Host plant attributes (leaf traits and phylogeny) were more important in determining the abundance of hub ASVs in temperate regions, while spatial-related effects were more important in influencing tropical hub ASVs (Fig. 6). Additionally, we found a strong phylogenetic signal in the responses of ASVs to habitat covariance, indicating that phylogenetically closely related ASVs have similar habitat preferences (Fig. S8).

**Figure 5.**
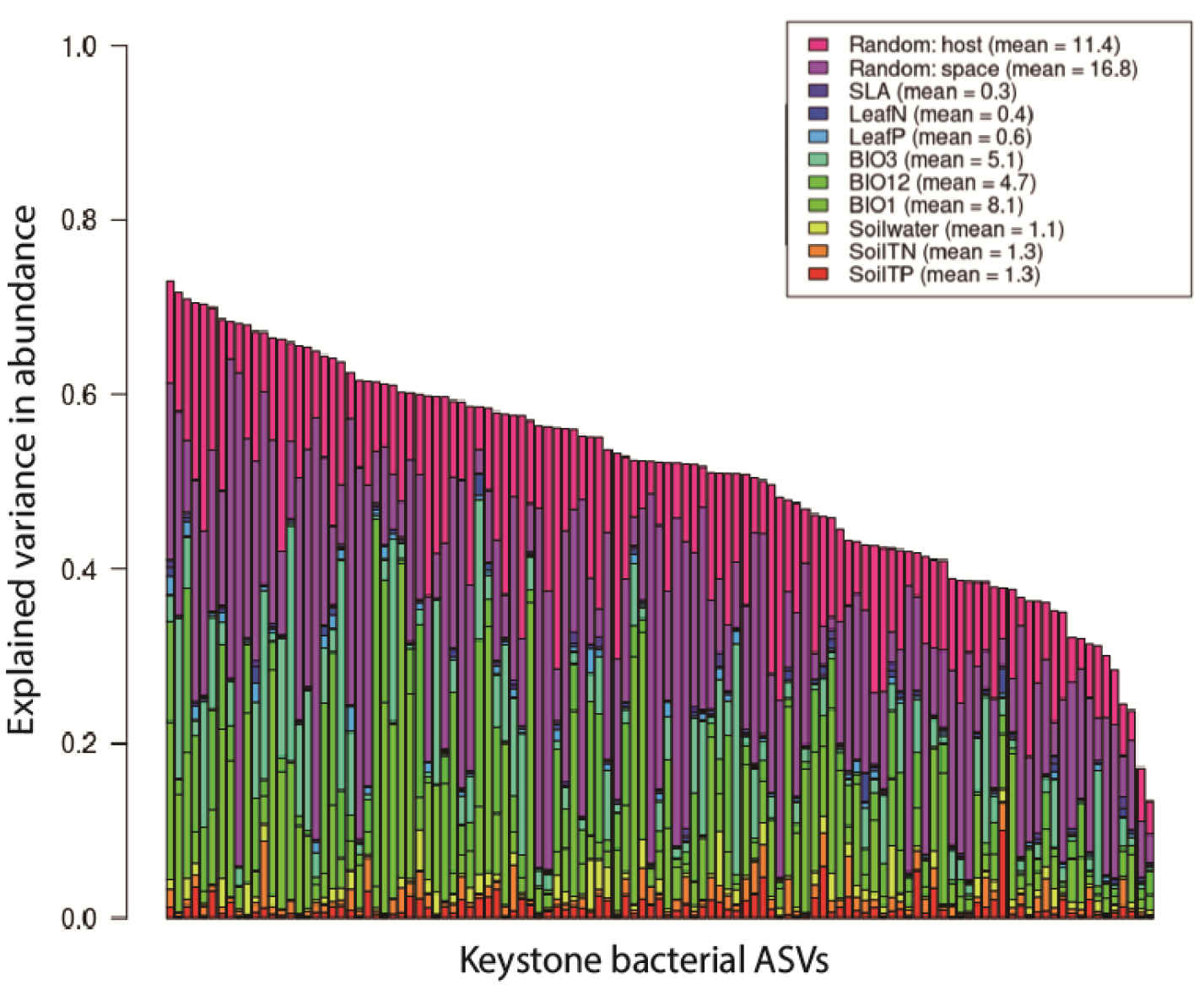
The results of joint species distribution modeling, showing (a) the absolute variation in bacterial abundance explained by each factor, the relative contribution of (b) host plant traits and (c) spatial factors on determining the abundance of ASVs in different regions. 115 ASVs were included in the model, and 9 habitat covariates and two distance matrices representing spatial and host phylogenetic relationship among samples were used as fixed and random term respectively. Each bar represents a hub bacterial ASV sorted according to the total variance explained, and the caption above the bar plot shows the averaged variation explained by each factor.

**Figure 6.**
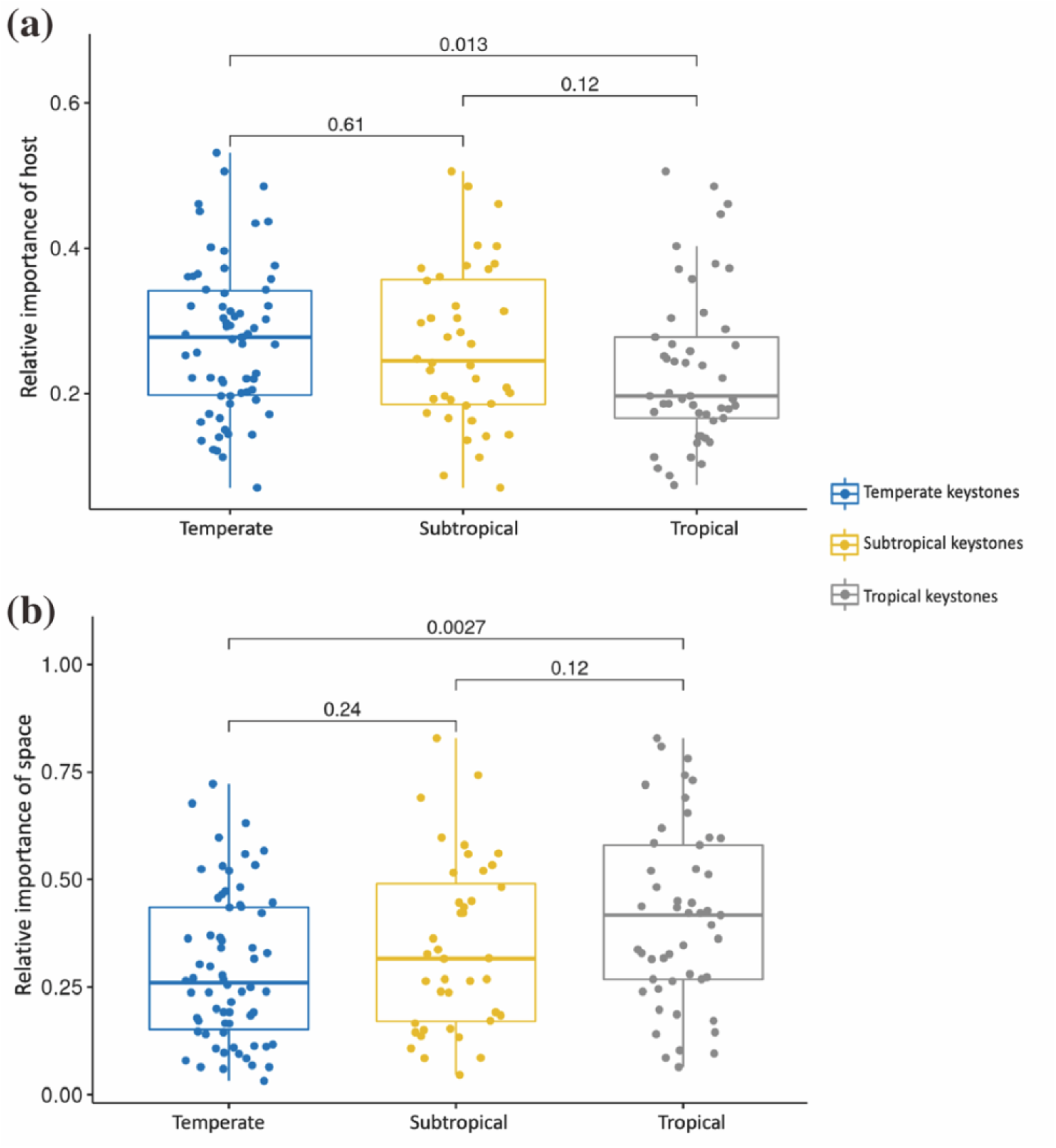
The contribution of host attributes (a) and spatial-related factors (b) to the abundance of hub taxa varies among regions. Numbers above the paired boxes symbolize the p values of pairwise Wilcoxin tests.

## Discussion

In this study, we revealed for the first time a strong latitudinal pattern in the diversity, community composition and distribution of plant-leaf associated bacteria, highlighting the varying strength of community assembly processes along the latitudinal gradient in shaping plant leaf-microbial symbioses. It is remarkable that we found a shift from strong host filtering in bacterial community assembly in temperate forests, to weak host filtering but a strong joint effect of multiple drivers in the tropics. These results suggest that plant-associated bacteria on leaves are formed following different rules across latitudes, with an increased complexity of different processes acting to explain bacterial symbioses at lower latitudes. The latitude-dependent community assembly processes were consistently evident in driving bacterial ASV richness, community composition and the abundance of metacommunity-level hub bacteria taxa, which together provide the most comprehensive understanding to date of the patterns and mechanisms of plant-bacterial associations at large geographical scales. Our findings imply responses of plant-bacteria symbioses to host plant composition and environmental change differ in tropical vs. temperate regions, serving as a baseline for predicting and managing plant-microbe symbioses under global change.

Although host attributes, environment, climate, plant neighborhoods and spatial factors have been shown to influence the diversity of phyllosphere bacteria in previous studies ^14,16,18–20^, most of these factors were analyzed individually in these studies. As such, their relative importance was unknown, which made it difficult to infer the processes underlying the observed host-microbe associations^31^. By simultaneously evaluating a broad range of factors across spatial scales, we have shown that host-related factors including plant phylogeny and leaf traits were the most important drivers of phyllosphere bacterial community assembly, especially in temperate regions. This result suggested that phylogenetically conserved host attributes including specific leaf area and leaf nutrient concentrations could select for leaf-associated bacteria^14,15^. Our results of independent effects of hosts on bacterial diversity at large spatial scales indicated consistent matching between host plant and phyllosphere bacteria regardless of climates, which represents a potential consequence of coevolutionary feedbacks between plant hosts and their associated microbes^15,32^. The importance of host plant selection mediated by phylogenetically conserved traits could further lead to host specialization of plant-associated microbes, which may allow microbes to coexist and explain their incredible diversity on leaves, and also contribute to the host-specific effects of plant-associated microbes in structuring plant communities^33–35^.

While host selection was among the most important factor in explaining phyllosphere bacterial community assembly, the joint effect of multiple other factors also played a critical role particularly in tropical forests. Many of these factors including spatial structure and neighboring plant communities are likely related to the importance of stochastic processes including dispersal for community assembly of plant-associated bacteria, suggesting a latitudinal patterns in the relative contribution of deterministic and stochastic process to shaping plant-bacteria association. This result concurred with the prediction of the niche-neutral continuum hypothesis that the importance of niche-based processes such as selection are stronger in low diversity temperate regions while the importance of species-neutral processes such as dispersal are more important in high diversity tropical regions ^22,36^. Previous studies have shown that the processes structuring plant community assembly increasingly deviated from neutrality with latitude, and here we provided the first evidence that this hypothesis was supported for microbes, suggesting that ecological hypotheses developed based on evidence from macroorganisms also apply to explain differences in microbial community assembly along latitudinal gradients^22^.

Although the composition of phyllosphere bacterial communities varied greatly across study sites along the latitudinal gradient, the different bacterial communities were interconnected by a small proportion of hub bacterial ASVs to form large-scale plant-bacterial cooccurrence networks. These hub ASVs were associated with a wide range of plant species in a broad geographic space and include taxa such as *Methylorubrum* and *Sphingomonas* that have been shown to play key roles in promoting plant health and productivity^10^. Additionally, the hub bacterial taxa at the metacommunity level could affect large-scale plant-bacteria interactions: the movement of hub taxa across local communities could mediate otherwise discrete ecological processes within a metacommunity. For example, although plant species differing in growth-mortality strategies hosted distinct leaf bacteria communities, the bacteria on one plant species can affect that of other plant species through their influence on the metacommunity of shared hub bacteria taxa^14,26^. Likewise, hub taxa could synchronize coevolutionary processes and control the pace of host-bacteria coevolution at metacommunity scales through gene flow^37^.

Thus, the spatial distribution of hub bacteria taxa should have a major impact on large-scale patterns of plant-bacteria associations^26^. The distribution of hub bacterial ASVs was clustered along the latitudinal gradient and was mainly controlled by climate, host attributes and spatial factors. This suggests that climate change and plant species range shifts could change plant-bacteria interactions through affecting the distribution of those hub taxa. Moreover, the influence of host plant attributes on bacterial abundance was more important for hub taxa in temperate than that in tropical forests, while spatial-related factors were more important for tropical than temperate hub taxa, consistent with the observation of varying strengths of different processes structuring bacterial diversity and community assembly along the latitudinal gradient. This result emphasizes the importance of host selection in temperate regions and dispersal in the tropics to explain the spatial distribution of hub bacteria taxa.

Understanding the response of plant microbiomes to global change and the consequent impact on plant communities and ecosystem function requires large-scale studies on the diversity and biogeography of plant-associated microbes^7,30,38^. Our results show for the first time that the community assembly of plant leaf-associated bacterial communities is driven by processes whose importance varies along a latitudinal gradient. A corollary of this finding is that global change is likely to have different effects on plant leaf-microbe associations along the latitudinal gradient, with tropical associations being more sensitive to climate warming and environmental changes and temperate associations being more sensitive to changes in host plant abundance and plant species range shifts. The direction of the global change effect could even reverse from negative to positive from low latitudes to high latitudes, as recently shown for plant soil microbes^39,40^. Host selection could result in distinct microbiome compositions that exert plant species-specific impacts on host fitness. Although our study showed varying strength of host selection across leaf microbiomes from temperate to tropical forests, it remains to be understood whether the impact of plant leaf microbiomes on plant communities differs along latitude gradients. Additional broad-scale comparative studies that take multiple factors into account will be required to reveal the role of plant microbiomes in mediating the impact of global change on plant communities and ecosystem function.

## Methods

### Study sites

We conducted the study at 10 forest sites across China with latitude ranging from 18.7°N to 51.8°N (Table S2). The sites are part of a global network of stem-mapped forest plots of the Forest Global Earth Observatory (ForestGEO http://forestgeo.si.edu/). For each site, we selected approximately 1/3 of the tree species and around 3 individuals per species. The choice of these species considered both shade tolerant and intolerant, early-and late-successional species, in order to have a gradient in host attributes that may explain the variation in bacterial community assemblages. In total, 1453 trees from 328 plant species of 148 genera and 59 families were sampled.

### Sampling and processing

For each tree individual, 50-100g leaves from the sub-canopy (2-10 m above ground) were collected. Leaves were placed in a sterile plastic bag and washed using 100 mL of a 1:50 dilution of Redford buffer (1M Tris, 500 mM EDTA, 1.2% CTAB) within 24 hours after sampling^14^. The suspension was collected and centrifuged at 3300 g for 25 minutes and the pellet was added into an extraction tube to store in -20 °C until DNA extraction. Microbial DNA was extracted following the manufacturer’s protocol (QIAGEN Powersoil DNA extraction kit) and the V5-V6 region of bacteria 16S rRNA genes was amplified using primers 799F and 1115R to determine the composition of the phyllosphere bacterial community. PCR products were normalized using a SequalPrep Normalization kit (Thermo Fisher Scientific), pooled and then purified using AMPure to remove contaminants. The DNA library was prepared by mixing equimolar concentrations of DNA from each sample, and then sequenced using Illumina MiSeq Reagent Kit v3 for 300 bp paired-end sequencing.

### Bioinformatics

We used DADA2 to identify the amplicon sequence variants (ASV) observed in each sample^41^. The base quality scores (Q) were calculated, and the nucleotides at 19-240 bp and 16-240 bp positions were kept for forward and reverse DNA sequences respectively. Then we inferred error rates and inferred ASVs using DADA2 default parameters^41^. Paired read ends were merged with a minimum overlap of 20. Non-target sequences and chimeras were removed. Taxonomy was assigned to each ASV by comparison with the SILVA SSU r138 database.

We revealed around 116 million high-quality 16S rRNA gene sequences and identified 55693 amplicon sequence variants (ASVs) from 1453 samples. Sequence abundance per sample ranged from 15485 to 522574, which could bias estimates of ASV diversity (Fig. S9a). Thus, we randomly rarefied the ASV table to 15000 sequences per sample using the R package ‘vegan’, and 49254 ASVs were retained after rarefaction. This eliminated only a small proportion of extremely rare ASVs and had little impact on beta diversity (e.g. community dissimilarity; Fig. S9b). Most ASVs (82%) were identified to the family level, 58% were classified to genus and only 4% were identified to the species level.

### Metadata

We collected metadata describing host plant attributes, abiotic environments, neighborhood plants and spatial location. Plant phylogenetic tree was derived from a phylogeny database for vascular plants according to the species identity using the *V*.*phyloMaker* function in R ^42^. Host plant identity and functional traits and abiotic environments including topographic and edaphic variables were derived from the base data of each site. Climate variables including annual mean temperature, annual precipitation, and precipitation seasonality were collected from WorldClim database based on the geographical coordinates of sites. We calculated the richness, abundance, mean phylogenetic distance and mean nearest neighbor distance of neighborhood plants present within 10 meters around the sample, and included the local abundance of host plant species in the metadata (Table S3).

### Statistical analysis

We used R (version 3.6) for all statistical analyses. The diversity of phyllosphere bacteria was modelled as a function of variables of host, environment, neighborhood plants and spatial factors. To prepare these variables, Moran’s Eigenvector Maps (MEM) was used to decompose geographic distance matrices into a series of orthogonal eigenvectors representing the spatial relationships among samples, and principal coordinates of neighbor matrices (PCNM) was applied on the phylogenetic distance matrix among host plants to generate a series of variables that representing the position of host plant in a multi-dimensional phylogenetical space. Both phylogenetic and spatial eigenvectors can be used in regression to take phylogenetic/spatial autocorrelation into account ^43^. Environment, host traits and diversity of plant neighborhoods variables were rescaled using the formula (*x* – *x*_*min*_)/(*x*_*max*_ – *x*_*min*_) to standardize the effects of the variables. Pairwise Pearson’s correlation coefficients were calculated to test the correlation among all variables, and for predictors with correlation coefficient > 0.9, one predictor was randomly removed to reduce multicollinearity.

Species richness was calculated as the number of ASVs with abundance > 1, which follows a negative binomial distribution (NBD). To model the effects of variables on richness, we used generalized linear model with NBD as the error structure, implemented with function *glm*.*nb* in the R package MASS. We first built a candidate set of generalized linear models to represent the individual and joint linear contributions for types of mechanisms (host, environment, neighborhoods and spatial proximity) using the predictors for each mechanism, resulting in a total of 15 models. For each model, a stepwise selection procedure was used for model selection based on Akaike’s information criterion corrected for small sample size (AICc). Non-significant variables (P < 0.05) were further excluded using the *update* function. Variance inflation factor (VIF) was calculated and variables with VIF > 5 were also removed to avoid multicollinearity. To quantify the individual effects of four types of variables and their interactions on bacterial richness, we used variation partitioning where the independent effect of each factor was evaluated as the difference in *R*^2^ between models with and without this factor, and the interactive effect was calculated by subtracting the independent effect from overall effects. We used a subset of samples to evaluate the effect of host plant traits as the trait data are available for most but not all samples (Table S3). The effect of plant traits was highly corrected with that of plant phylogeny (Fig. S10). Thus, we only included plant phylogenetic eigenvectors as host variables in the variation partitioning analysis.

We repeated the same variance partition analysis on bacterial composition. To do that, the ASV table was Hellinger transformed to represent the community composition of bacteria. We used redundancy analysis (RDA) to model the effect of those variables on community composition. Variable selection was done for each model using forward selection implemented with the function *forward*.*sel* in R package packfor following the recommended stopping rules, and variation partitioning was implemented by *varpart* function in the ‘vegan’ package ^44^.

The analyses of richness and community composition were conducted at local scales for the 10 sites, at regional scales for tropical, subtropical and temperate forests, and at the global scale by combining all the sites. Specifically, Heishiding (HSD) and Dinghushan (DHS) sites (located at 23.27°N, 23.17°N respectively) were on the northern boundary of the tropics.

Although classified as subtropical sites, the two forests are more similar in vegetation composition and climate conditions with the tropical forest Jianfengling (JFL, 18.73°N) than those typical subtropical forests (Table S6). We thus included HSD and DHS as ‘tropical’ sites, resulting in 3 tropical, 4 subtropical and 3 temperate sites. The relative importance of each factor and their interactive effects were evaluated as the proportion of explained variance, which was further related to latitude using a linear regression.

To construct networks that represent plant-bacteria associations at different geographical scales, we first transformed ASV abundances to presence-absence with a threshold of 15 reads per sample (0.1% of the total read count of each sample) to be considered present, in order to minimize the effects of extreme rare species and sequencing errors on network construction. The occurrence of ASV in samples was then converted to occurrences on plant species according to the plant host identity of each sample, where a cell entry represented the number of leaf samples that were observed to have the ASV occurring on a plant species. In this step, the same plant species from different sites were represented in different rows as we were interested in how metacommunity bacteria interconnect local plant-bacteria networks across sites. The species-ASV table comprises 490 rows and 12065 columns, and the binary (presence/absence) data of this table were used to construct the network. We constructed 10 local networks representing plant-bacteria associations at site level, 3 regional networks for temperate, subtropical and tropical regions, and a global network using ‘igraph’ package in R. The regional and global networks were visualized using program GePhi with the ‘ForceA-tlas2’ layout algorithm, where plant species from different sites were highlighted using warm colors. For these networks, we identified hub ASVs in core positions of the networks by calculating the betweenness centrality score of each node/ASV. Betweenness centrality was standardized using the formula (*x* – *x*_*min*_)/(*x*_*max*_ – *x*_*min*_) to make it range from 0 to 1, and ASVs with scaled betweenness above 0.5 were recognized as hubs^26^. A total of 115 ASVs were identified as metacommunity-level hubs at regional and/or global scales.

The biogeography of the hub ASVs was first explored using a heatmap that mapped the frequency of ASVs (the proportion of samples in which the ASVs occurred) for each site. We then applied a joint species distribution modeling approach, called Hierarchical Modelling of Species Composition (HMSC)^45^, to predict the abundance of the 115 hub ASVs at the global scale. We selected a set of habitat variables that indicate important mechanisms potentially structuring bacterial community assembly, including 3 soil variables (total nitrogen, total phosphorus and water content), 3 climate variables (temperature, precipitation and isothermality), and 3 plant leaf traits (nitrogen, phosphorus and specific leaf area). As plant trait data were not available for all samples, 1242 out of 1453 samples with available plant trait data were used in this analysis. The 9 variables were treated as fixed terms to evaluate the effect of habitat conditions on ASVs, and two distance matrices representing the spatial and host phylogenetic relationships between samples were used as random terms to control for the effects of spatially and phylogenetically clustered data.

We applied a hurdle model, i.e., one model for presence-absence and another one for abundance conditional on presence, in order to quantify the drivers of both presence-absence and abundance of hub ASVs. We applied probit regression in the presence-absence model, and linear regression for transformed count data in the abundance model. The count data were transformed by declaring zeros as missing data, log-transforming, and then scaling the data to zero mean and unit variance within each species^45^. We fitted the HMSC model with the R-package Hmsc assuming the default prior distributions. We sampled the posterior distribution with five Markov Chain Monte Carlo (MCMC) chains, each of which was run for 37,500 iterations, of which the first 12,500 were removed as burn-in. We examined MCMC convergence by examining the potential scale reduction factors of the model parameters. The MCMC convergence of our model was satisfactory: the potential scale reduction factors for the β-parameters (that measures the responses of the species to environmental covariates) were on average 1.013 and 1.04 for abundance and presence-absence model respectively. To compare the species distribution patterns of ASV among regions, we calculated the relative importance of each mechanism: the fixed effects of host traits, soil condition and climate, and the random effects of space and host phylogeny, which were compared among temperate, subtropical and tropical hubs using paired samples Wilcoxon tests.

## Author contribution

Z.W., F.H. and S.W.K. conceived the project; Z.W. conducted field sampling and phyllosphere data collection; Y.J., M.Z., C.C., Y.C., S.F., G.J., M.J., J.L., Y.L., Y.L., K.M., X.M., X.Q., X.W., X.W., H.X., W.Y., L.Z., Y.Z. collected and collated the metadata and assisted with fieldwork; Z.W. analyzed data with supported from S.W.K.; Z.W. drifted the manuscript with input from S.W.K. and F.H.; All authors contributed to editing of the manuscript.

## References

1. Gilbert, J. A. & Neufeld, J. D. Life in a world without microbes. Plos Biol 12, e1002020 (2014).

2. Berendsen, R. L., Pieterse, C. M. J. & Bakker, P. A. H. M. The rhizosphere microbiome and plant health. Trends Plant Sci 17, 478–486 (2012).

3. Liu, H., Brettell, L. E. & Singh, B. Linking the phyllosphere microbiome to plant health. Trends Plant Sci 25, 841–844 (2020).

4. Linde, S. van der et al. Environment and host as large-scale controls of ectomycorrhizal fungi. Nature 558, 243–248 (2018).

5. Martin, F. M., Uroz, S. & Barker, D. G. Ancestral alliances: Plant mutualistic symbioses with fungi and bacteria. Science 356, (2017).

6. Vorholt, J. A. Microbial life in the phyllosphere. Nat Rev Microbiol 10, 828–840 (2012).

7. Cavicchioli, R. et al. Scientists’ warning to humanity: microorganisms and climate change. Nat Rev Microbiol 17, 569–586 (2019).

8. Perreault, R. & Laforest-Lapointe, I. Plant-microbe interactions in the phyllosphere: facing challenges of the anthropocene. Isme J 16, 339–345 (2022).

9. Lindow, S. E. & Leveau, J. H. J. Phyllosphere microbiology. Curr Opin Biotech 13, 238–243 (2002).

10. Chen, T. et al. A plant genetic network for preventing dysbiosis in the phyllosphere. Nature 580, 653–657 (2020).

11. Zamioudis, C. & Pieterse, C. M. J. Modulation of host immunity by beneficial microbes. Mol Plant-microbe Interactions 25, 139–150 (2012).

12. Moyes, A. B. et al. Evidence for foliar endophytic nitrogen fixation in a widely distributed subalpine conifer. New Phytol 210, 657–668 (2016).

13. Vacher, C. et al. The phyllosphere: microbial jungle at the plant–climate interface. Annu Rev Ecol Evol Syst 47, 1–24 (2016).

14. Kembel, S. W. et al. Relationships between phyllosphere bacterial communities and plant functional traits in a neotropical forest. Proc National Acad Sci 111, 13715–13720 (2014).

15. Lajoie, G. & Kembel, S. W. Plant-bacteria associations are phylogenetically structured in the phyllosphere. Mol Ecol 30, 5572–5587 (2021).

16. Laforest-Lapointe, I., Messier, C. & Kembel, S. W. Host species identity, site and time drive temperate tree phyllosphere bacterial community structure. Microbiome 4, 27 (2016).

17. Laforest-Lapointe, I., Paquette, A., Messier, C. & Kembel, S. W. Leaf bacterial diversity mediates plant diversity and ecosystem function relationships. Nature 546, 145–147 (2017).

18. Lajoie, G. & Kembel, S. W. Host neighborhood shapes bacterial community assembly and specialization on tree species across a latitudinal gradient. Ecol Monogr (2021) doi:10.1002/ecm.1443.

19. Leducq, J.-B. et al. Fine-scale adaptations to environmental variation and growth strategies drive phyllosphere methylobacterium diversity. Mbio 13, e03175–21 (2022).

20. Finkel, O. M. et al. Distance-decay relationships partially determine diversity patterns of phyllosphere bacteria on tamrix trees across the sonoran desert. Appl Environ Microb 78, 6187– 6193 (2012).

21. Magan, N. & Baxter, E. Effect of increased CO 2 concentration and temperature on the phyllosphere mycoflora of winter wheat flag leaves during ripening. Ann Appl Biol 129, 189–195 (1996).

22. Qiao, X., Jabot, F., Tang, Z., Jiang, M. & Fang, J. A latitudinal gradient in tree community assembly processes evidenced in Chinese forests. Global Ecol Biogeogr 24, 314–323 (2015).

23. Dyer, L. A. et al. Host specificity of Lepidoptera in tropical and temperate forests. Nature 448, 696–699 (2007).

24. Banerjee, S., Schlaeppi, K. & Heijden, M. G. A. van der. Keystone taxa as drivers of microbiome structure and functioning. Nat Rev Microbiol 16, 567–576 (2018).

25. Agler, M. T. et al. Microbial hub taxa link host and abiotic factors to plant microbiome Variation. Plos Biol 14, e1002352 (2016).

26. Toju, H. et al. Species-rich networks and eco-evolutionary synthesis at the metacommunity level. Nat Ecol Evol 1, 0024 (2017).

27. Leibold, M. A. et al. The metacommunity concept: a framework for multi-scale community ecology. Ecol Lett 7, 601–613 (2004).

28. Thompson, J. N. The Geographic mosaic of coevolution. (2005) doi:10.7208/chicago/9780226118697.001.0001.

29. Toju, H., Tanabe, A. S. & Sato, H. Network hubs in root-associated fungal metacommunities. Microbiome 6, 116 (2018).

30. Martiny, J. B. H. et al. Microbial biogeography: putting microorganisms on the map. Nat Rev Microbiol 4, 102–112 (2006).

31. Hanson, C. A., Fuhrman, J. A., Horner-Devine, M. C. & Martiny, J. B. H. Beyond biogeographic patterns: processes shaping the microbial landscape. Nat Rev Microbiol 10, 497– 506 (2012).

32. Jr, P. R. G., Jordano, P. & Thompson, J. N. Evolution and coevolution in mutualistic networks. Ecol Lett 14, 877–885 (2011).

33. Wang, Z. et al. Effects of host phylogeny, habitat and spatial proximity on host specificity and diversity of pathogenic and mycorrhizal fungi in a subtropical forest. New Phytol 223, 462– 474 (2019).

34. Lajoie, G. & Parfrey, L. W. Beyond specialization: re-examining routes of host influence on symbiont evolution. Trends Ecol Evol 37, 590–598 (2022).

35. Semchenko, M. et al. Deciphering the role of specialist and generalist plant–microbial interactions as drivers of plant–soil feedback. New Phytol (2022) doi:10.1111/nph.18118.

36. Gravel, D., Canham, C. D., Beaudet, M. & Messier, C. Reconciling niche and neutrality: the continuum hypothesis. Ecol Lett 9, 399–409 (2006).

37. Urban, M. C. et al. The evolutionary ecology of metacommunities. Trends Ecol Evol 23, 311–317 (2008).

38. Brunel, C. et al. Towards unraveling macroecological patterns in rhizosphere microbiomes. Trends Plant Sci 25, 1017–1029 (2020).

39. Bachelot, B. et al. Altered climate leads to positive density-dependent feedbacks in a tropical wet forest. Global Change Biol 26, 3417–3428 (2020).

40. Liu, Y. & He, F. Warming intensifies soil pathogen negative feedback on a temperate tree. New Phytol 231, 2297–2307 (2021).

41. Callahan, B. J. et al. DADA2: High-resolution sample inference from Illumina amplicon data. Nat Methods 13, 581–583 (2016).

42. Jin, Y. & Qian, H. V. PhyloMaker: an R package that can generate very large phylogenies for vascular plants. Ecography 42, 1353–1359 (2019).

43. Dray, S., Legendre, P. & Peres-Neto, P. R. Spatial modelling: a comprehensive framework for principal coordinate analysis of neighbour matrices (PCNM). Ecol Model 196, 483–493 (2006).

44. Peres-Neto, P. R., Legendre, P., Dray, S. & Borcard, D. Variation partitioning of species data matrices: estimation and comparison of fractions. Ecology 87, 2614–2625 (2006).

45. Ovaskainen, O. et al. How to make more out of community data? A conceptual framework and its implementation as models and software. Ecol Lett 20, 561–576 (2017).

