## Supplemental Figs and Tables for "A latitudinal pattern of plant leaf-associated bacterial community assembly"

**The supplementary material includes:**

Figure S1 to S10
Tables S1 to S3

**Figure S1** Class-(a) and Order-(b) and Family-level(c) taxonomic composition of bacteria ASVs across the 10 sites. Only taxa with relative abundance above 0.1% are shown.

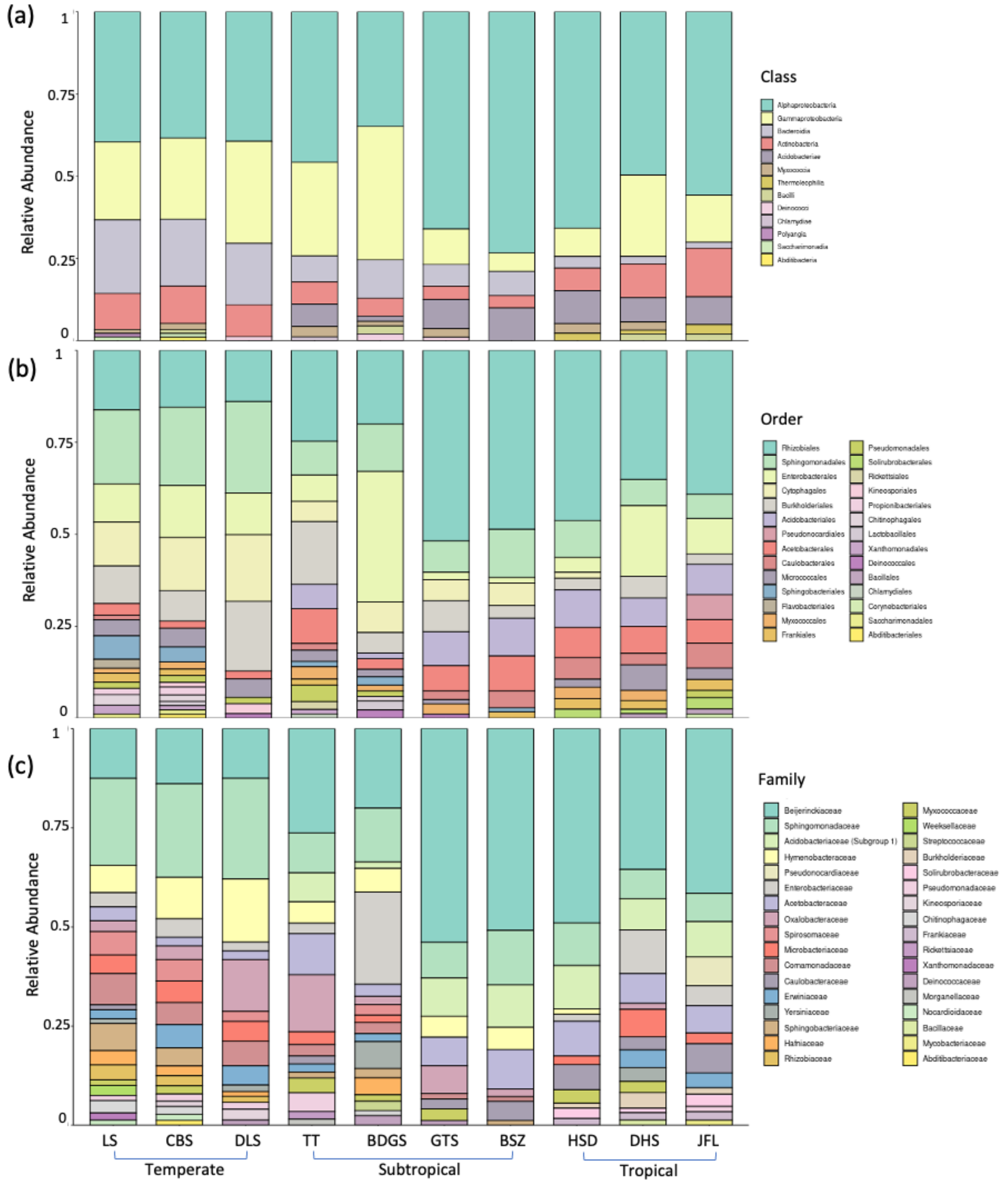

**Figure S2** Linear regressions on ASV richness of phyllosphere bacteria with (a) elevation, (b) slope, (c) annual mean temperature and (d) the richness of neighborhood plants. Second-order polynomial fits were shown in (a) and (c). Statistical p-values and the explanatory power ($R^2$) of models are reported.

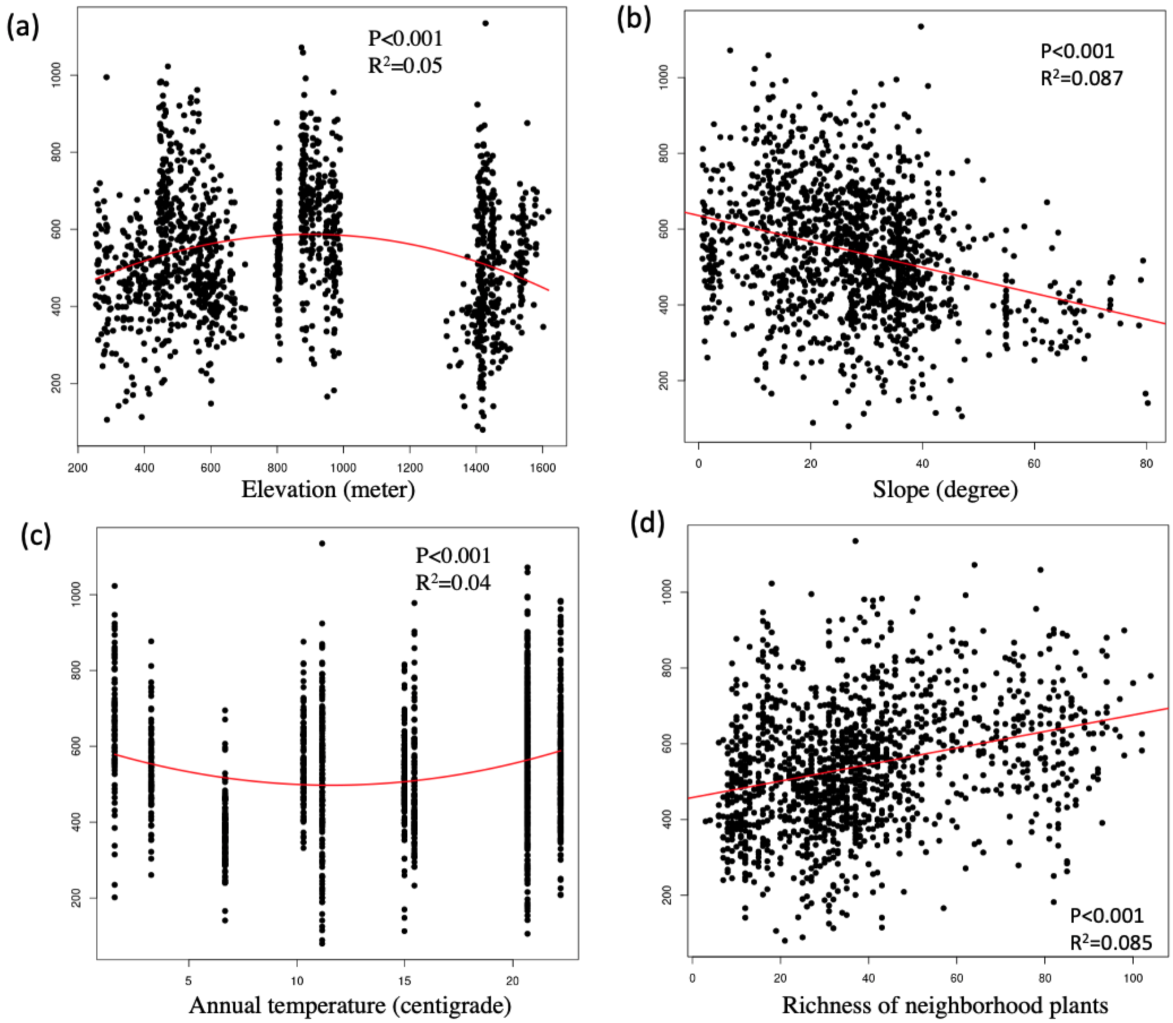

**Figure S3** Nonmetric multidimensional scaling (NMDS) ordination of variation in phyllosphere bacterial community structure on tree leaves. Ellipses represent 95% confidence intervals around samples from different host plant taxonomic families. The ordination was based on Bray-Curtis dissimilarity of Hellinger transformed community data. A total of 1016 samples from 17 plant families that have more than 30 samples were shown. Arrows inside plots indicate significant variables correlated with sample scores on each ordination axis and numbers in brackets show the explanatory power ($R^2$) of the correlation.

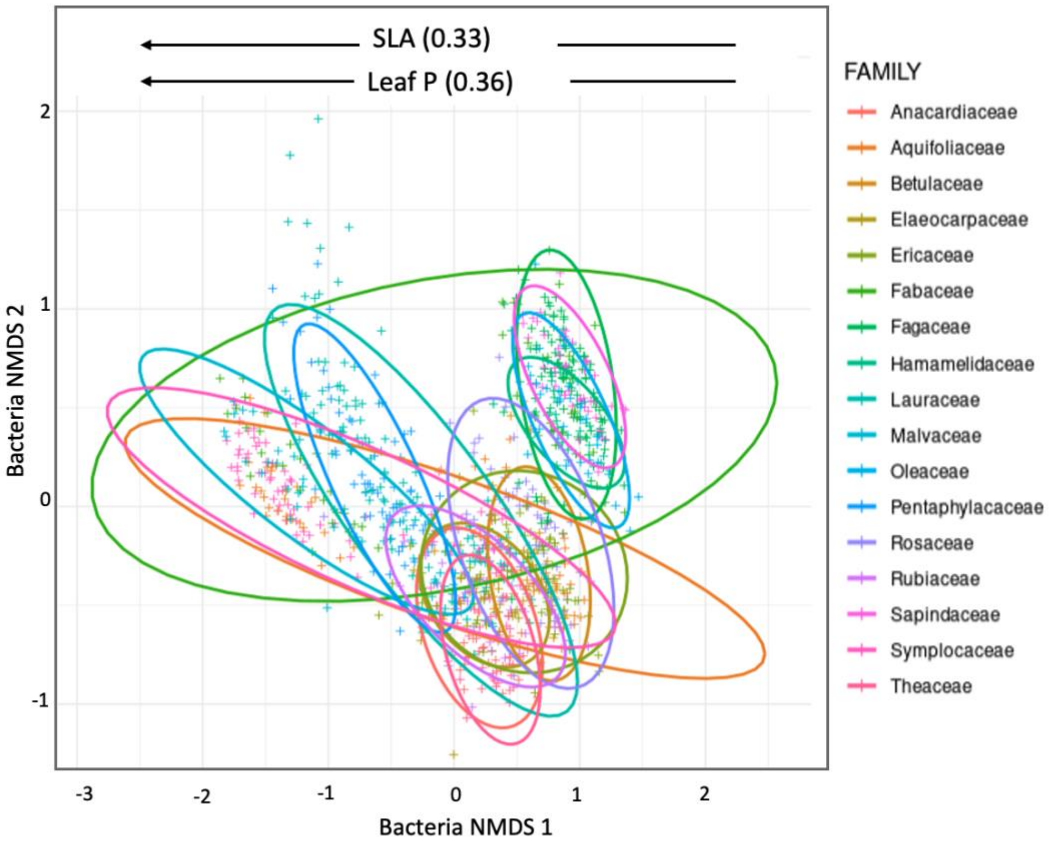

**Figure S4** Mantel correlation between bacterial communities at different geographic distance thresholds (a,c,e,g), and between bacteria community dissimilarity and phylogenetic distance of hosts (b,d,f,h). We show the patterns at local scales for a temperate forest CBS (a,b), a subtropical forest TT (c,d) and a tropical forest JFL (e, f) and at a global scale (g, h). A positive value with solid point in (a,c,e,g) indicates evidence for positive autocorrelation of bacterial community at that distance class ($p<0.05$). Mantel test for the similarity between community dissimilarity and phylogenetic distance in (b,d,f,h) showed strong evidence for positive correlations of the two matrices ($p<0.01$) in all cases.

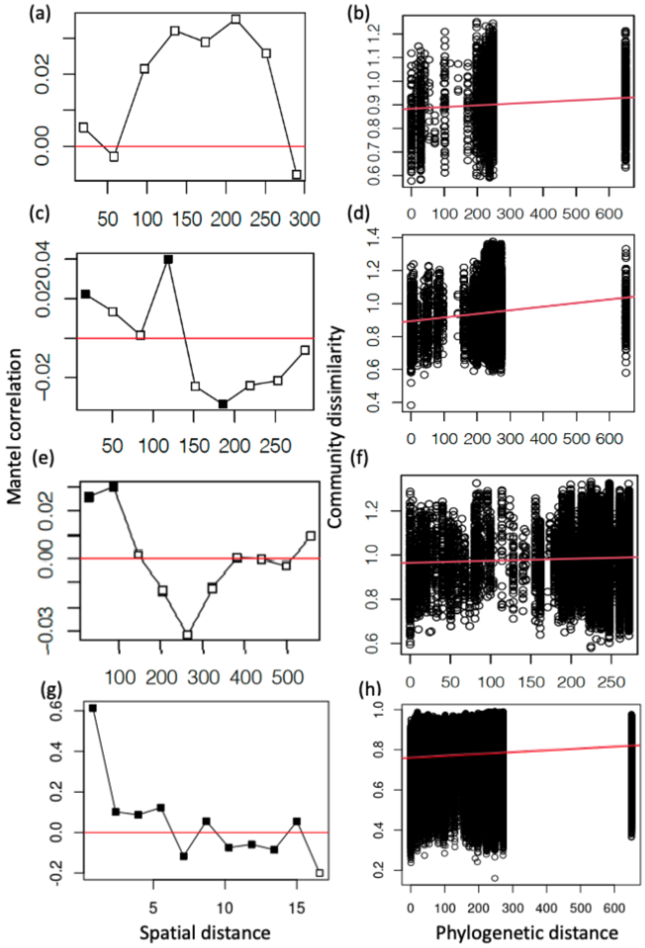

**Figure S5** Variation partitioning Venn diagrams showing the percentage of variation in species richness explained by host, abiotic environment, neighborhood plants and spatial factors at local scales for each site, regional scales for temperate, subtropical, and tropical sites, and a ‘global’ scale that includes all sites. The Venn diagrams show the absolute variance explained by each factor and the histograms on the right of Venn diagrams show the relative contribution of the independent and shared effects of four variables to the explained variance. Explained variation less than 1% was not shown in the Veen diagram. Abbreviation; Env: abiotic environments; Spa: spatial variables; Nei: neighborhood plant communities.

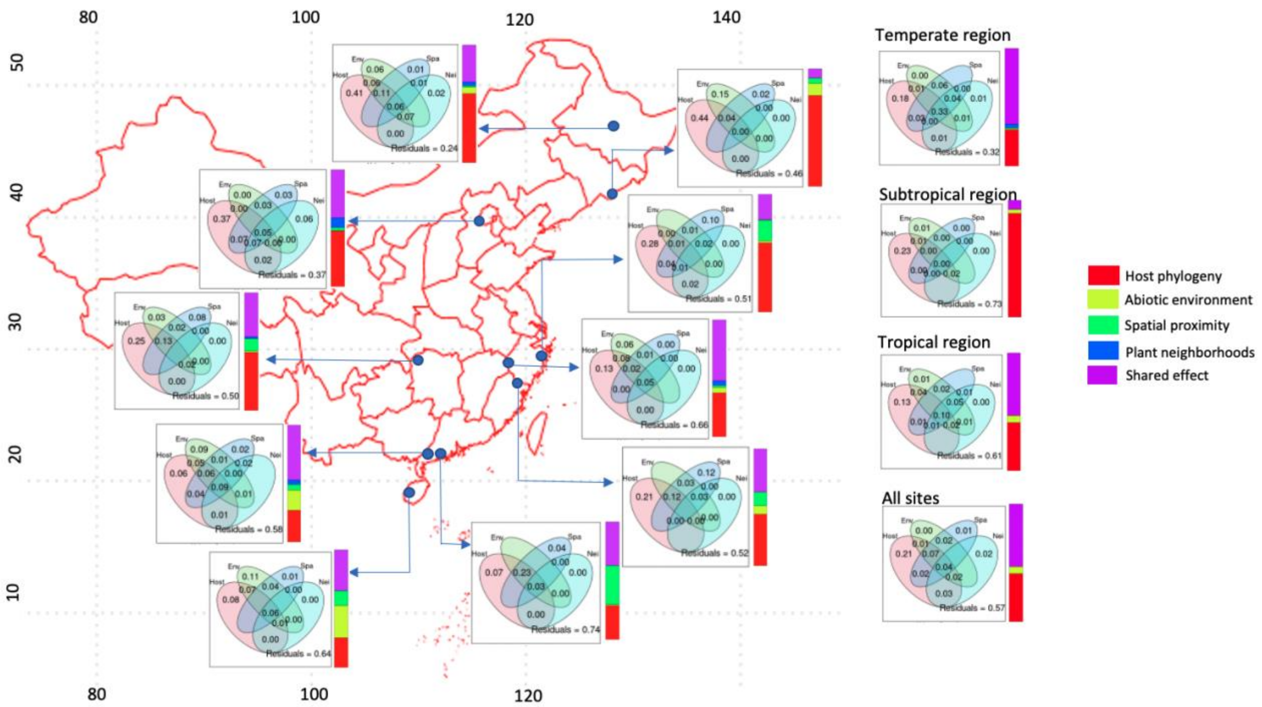

**Figure S6** Order-(a) and genus-level (b) taxonomic composition of the top 20 ASVs with the highest network betweenness centrality in plant-bacteria association networks. Local networks include 3 temperate (LS, CBS, DLS), 4 subtropical (TT, GTS, BDGS and BSZ) and 3 tropical forests (HSD, DHS, JFL), regional networks include temperate (Temp), Subtropical (Subtrop) and Tropical (Trop) regions, and a global-level network includes all sites.

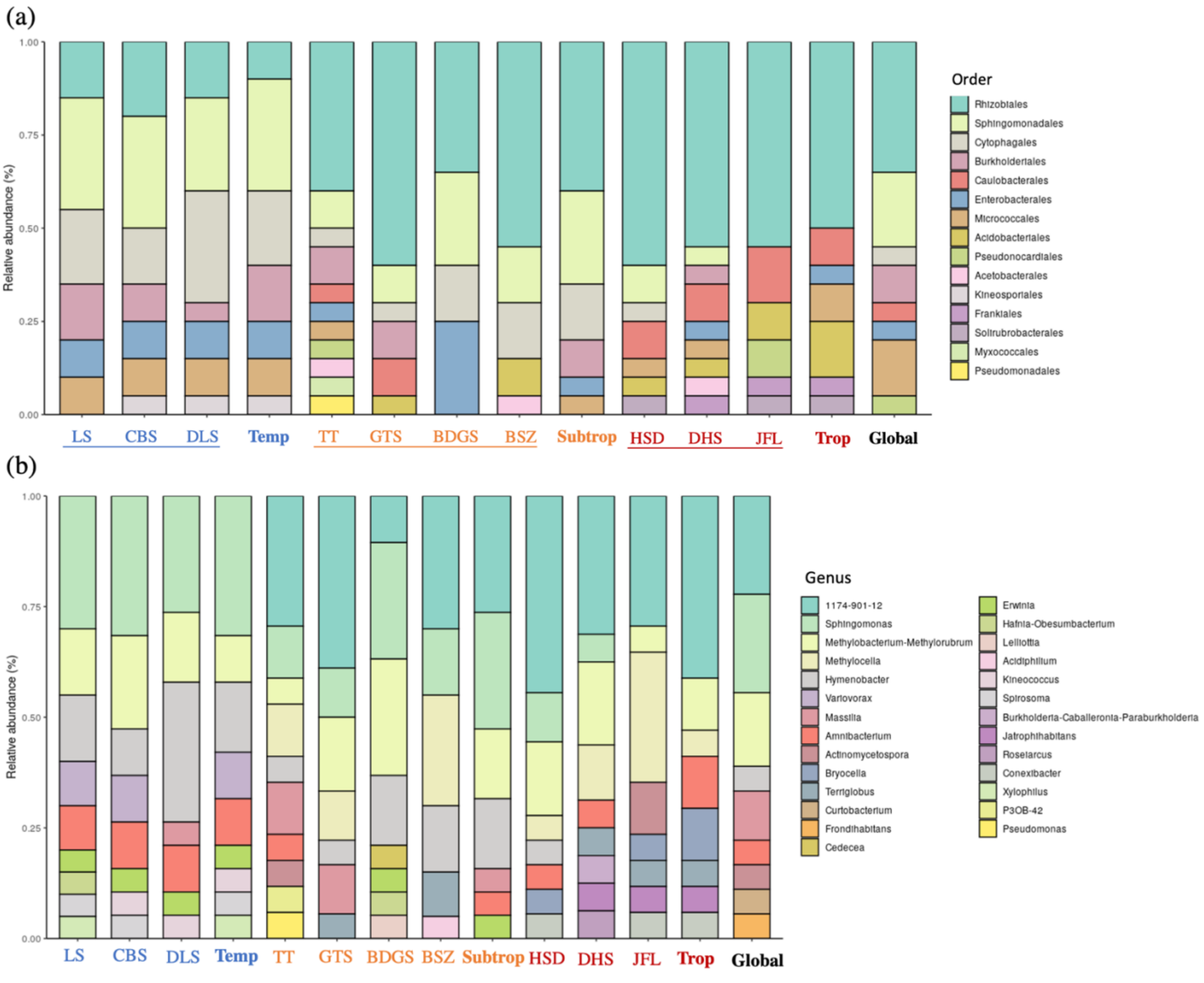

**Figure S7** The results of joint species distribution modeling, showing the absolute variation in bacterial presence-absence explained by different factors for each bacterial ASV. 115 ASVs were included in the model, and 9 habitat covariates and two distance matrices representing spatial and host phylogenetic relationship among samples were used as fixed and random term respectively. Each bar in represents a bacterial ASV sorted according to the total variance explained, and the caption above the barplot shows the average percent variation explained by each factor.

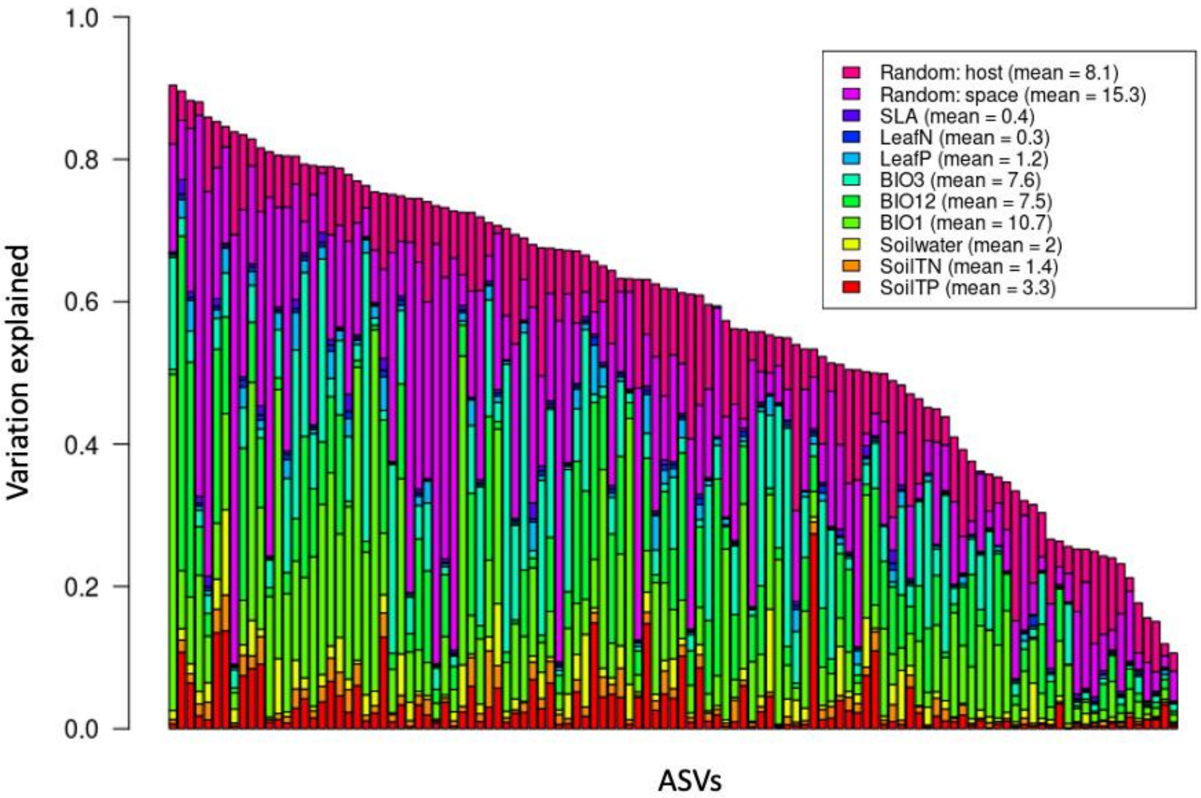

**Figure S8** The response of 115 hub ASVs to habitat covariates. Columns are habitat covariates and rows represent ASVs arranged according to their phylogeny. Red color indicates positive effects of habitat covariates on ASV abundance while blue indicate negative ones. Significant phylogenetic conservatism in the response was detected. Abbreviations, SoilTP: soil total phosphorus; SoilTN: soil total nitrogen; BIO1: annual temperature; BIO12: annual precipitation; BIO3: temperature isothermality; LeafP: plant leaf phosphorus content; LeafN: plant leaf nitrogen content; SLA: specific leaf area.

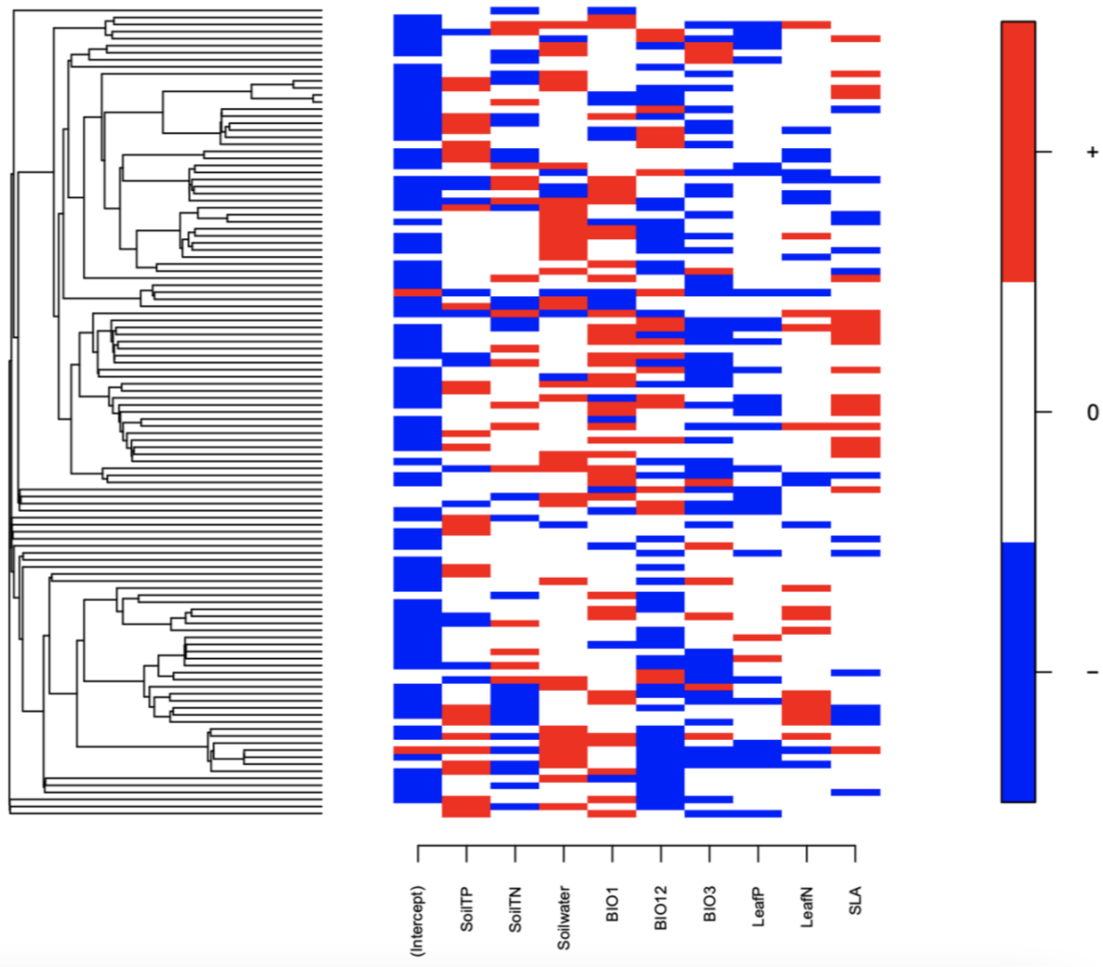

**Figure S9** The correlation between (a) sequencing depth and the species richness of amplicon sequence variants (ASVs), and (b) Hellinger-transformed Euclidean distance of community composition calculated using raw and rarefied ASV table.

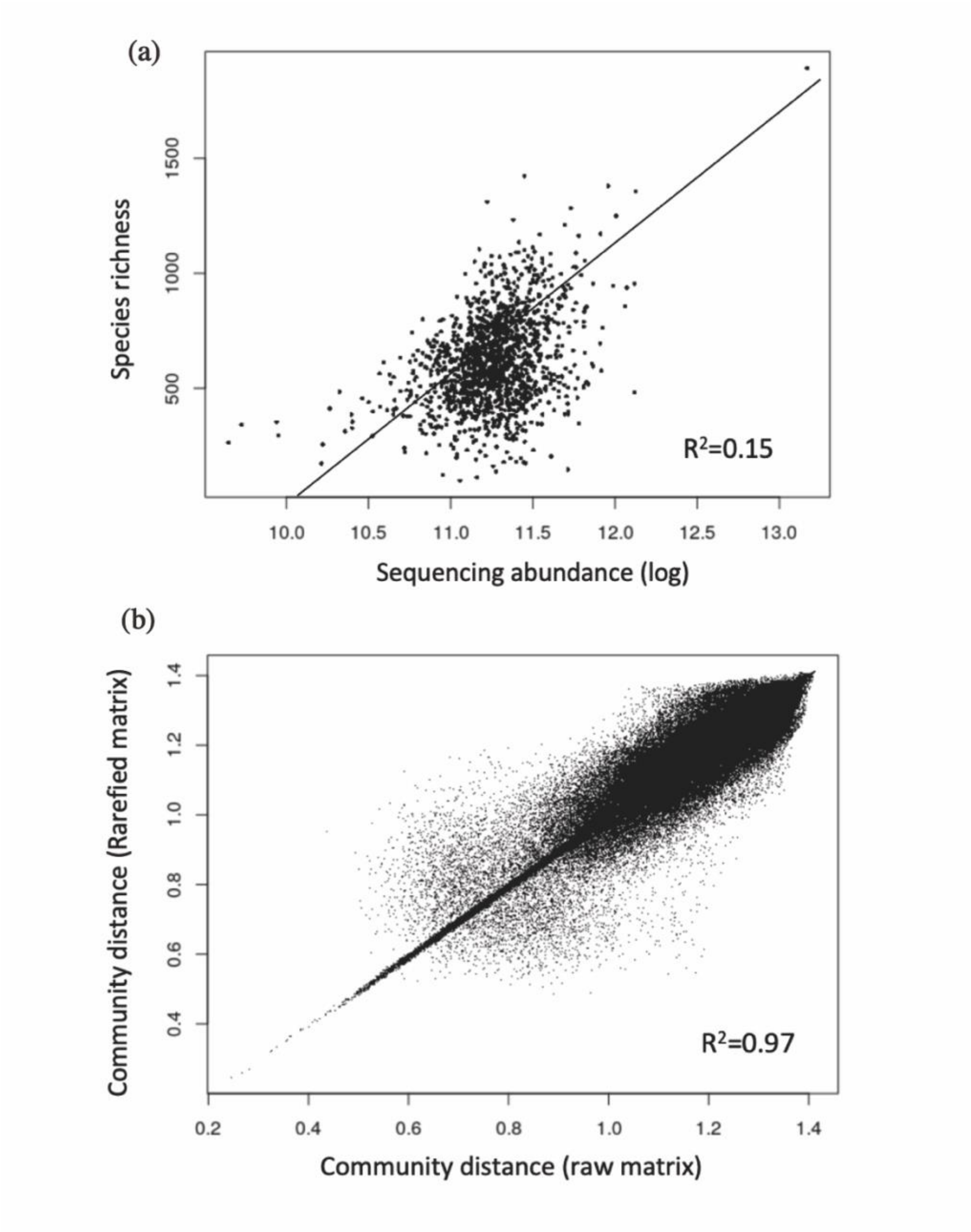

**Figure S10** Venn diagram showing the separate and confounding effect of host phylogeny and host traits on the species richness (a) and community composition (b) of phyllosphere bacteria for each site (site abbreviations are defined in Methods), regions and 'global'. We excluded site BSZ, site DLS and temperate region in this analysis as plant trait data are not available for most samples in these sites/regions.

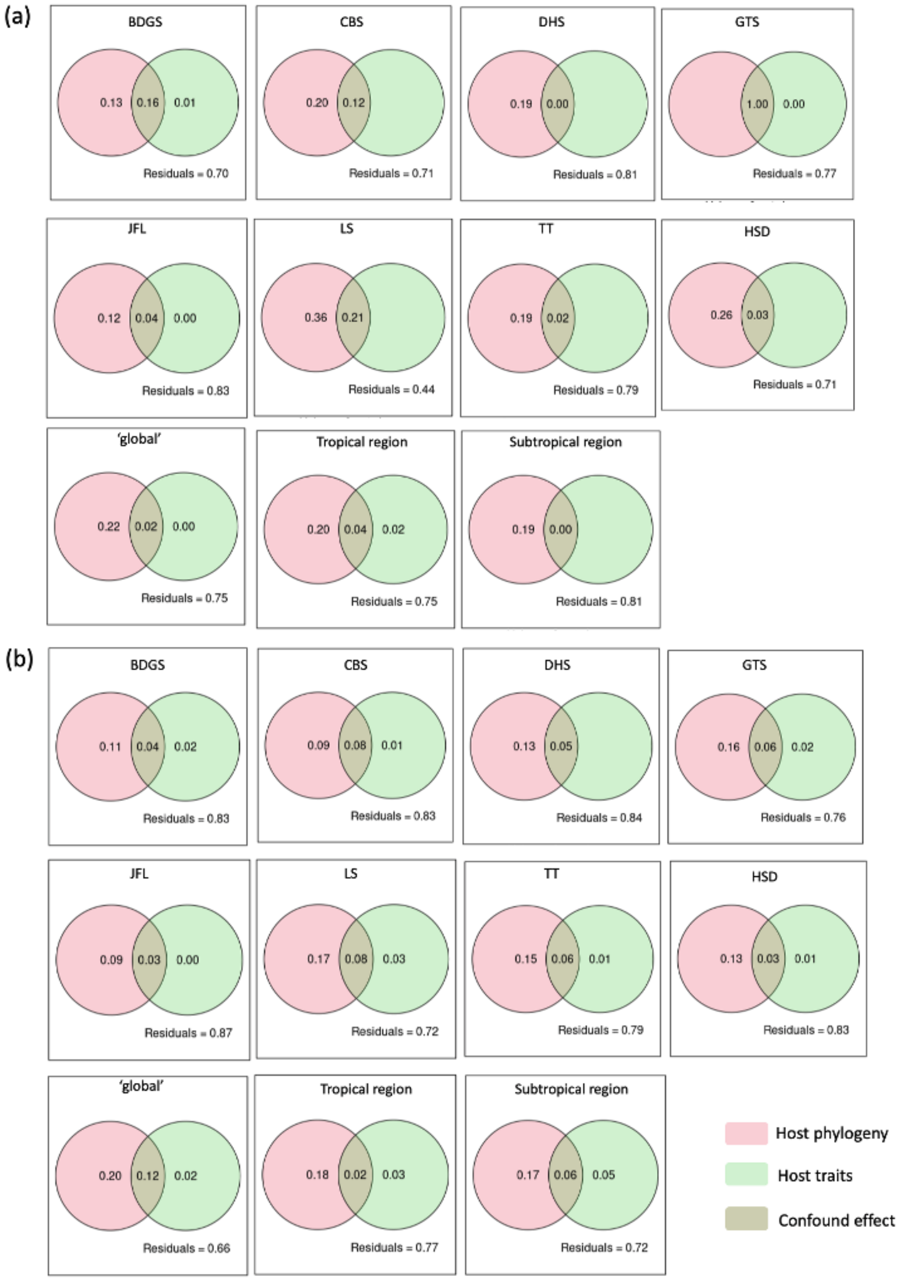

**Tables S1** The effect of abiotic environment (soil condition + topography, model1), climate variables (model2), neighborhood plants (model3) and host leaf traits (model4) on the community composition of phyllosphere bacteria. PERMANOVA analysis was applied on 1453 samples collected from 10 sites. We treated plant family identity as a random block in model 1 and 2, and site identity as a block in model 3 and 4. Abbreviations, SoilT*: total content in soil; SoilA*: available content; BIO1: annul mean temperature; BIO3: temperature isothermality; BIO12: Annual precipitation: BIO15: precipitation seasonality; aiet0: aridity; et0: evapotranspiration; richness.10, abundance.10, sesmntd.10: the richness, abundance and standardized effect size of mean nearest taxon distances of plant communities within 10 meters; SLA: specific leaf area.

| Variables | df | SS | $R^2$ | F | Pr(>F) | |
|---|---|---|---|---|---|---|
| **Model 1** | | | | | | |
| Elevation | 1 | 27.74 | 0.06014 | 139.5513 | 0.001 | |
| Slope | 1 | 12.3 | 0.02667 | 61.8976 | 0.001 | |
| SoilTN | 1 | 51.21 | 0.11103 | 257.6664 | 0.001 | |
| SoilTP | 1 | 32.08 | 0.06955 | 161.3968 | 0.001 | |
| SoilAN | 1 | 16.19 | 0.03511 | 81.4793 | 0.001 | |
| SoilAK | 1 | 5.19 | 0.01126 | 26.119 | 0.001 | |
| Soilwater | 1 | 11.61 | 0.02518 | 58.425 | 0.001 | |
| SoilpH | 1 | 11.05 | 0.02395 | 55.581 | 0.001 | |
| SoilOrgan | 1 | 5.33 | 0.01155 | 26.8045 | 0.001 | |
| Residual | 1442 | 286.61 | 0.62139 | | | |
| Total | 1452 | 461.25 | 1 | | | |
| **Model 2** | | | | | | |
| BIO1 | 1 | 81.72 | 0.17718 | 421.552 | 0.001 | *** |
| BIO3 | 1 | 33.5 | 0.07263 | 172.818 | 0.001 | *** |

| | | | | | | |
|---|---|---|---|---|---|---|
| BIO12 | 1 | 31.02 | 0.06726 | 160.02 | 0.001 | *** |
| BIO15 | 1 | 13.19 | 0.0286 | 68.057 | 0.001 | *** |
| aiet0 | 1 | 10.1 | 0.0219 | 52.116 | 0.001 | *** |
| et0 | 1 | 11.38 | 0.02467 | 58.702 | 0.001 | *** |
| Residual | 1446 | 299.64 | 0.63141 | | | |
| Total | 1452 | 474.55 | 1 | | | |
| **Model 3** | | | | | | |
| richness.10 | 1 | 60.61 | 0.1314 | 232.4606 | 0.001 | *** |
| abund.10 | 1 | 17.82 | 0.03863 | 68.3341 | 0.001 | *** |
| Sesmntd.10 | 1 | 3.81 | 0.00826 | 14.6118 | 0.001 | *** |
| Residual | 1448 | 377.52 | 0.81847 | | | |
| Total | 1452 | 461.25 | 1 | | | |
| **Model 4** | | | | | | |
| SLA | 1 | 27.27 | 0.07052 | 96.601 | 0.001 | *** |
| LeafN | 1 | 5.68 | 0.01469 | 20.116 | 0.451 | |
| LeafP | 1 | 4.26 | 0.01102 | 15.091 | 0.002 | ** |
| Residual | 1238 | 349.46 | 0.90378 | | | |
| Total | 1241 | 386.67 | 1 | | | |

**Table S2** Information on study sites, including the geographic coordinates, the size of the stem-mapped plot, the total number of stem and species in the plot, and the number of leaf samples taken in the study.

| Site | Coordinates | Size (ha) | No. Trees | No. species | No. Leaf Samples |
|---|---|---|---|---|---|
| Jianfengling | 18.7°N,108.9°E | 60 | 439676 | 289 | 299 |
| Heishiding | 23.3°N,111.5°E | 50 | 38004 | 236 | 190 |
| Dinghushan | 23.2°N,112.5°E | 20 | 71617 | 210 | 119 |
| Baishanzu | 27.7°N,119.2°E | 25 | 30945 | 132 | 117 |
| Gutianshan | 29.3°N,118.1°E | 24 | 140700 | 159 | 122 |
| Tiantongshan | 29.8°N,121.8°E | 20 | 115815 | 153 | 163 |
| Badagongshan | 29.7°N,110.0°E | 25 | 186575 | 232 | 149 |
| Donglingshan | 40°N,115.4°E | 20 | 52136 | 58 | 120 |
| Changbaishan | 42.4°N,128.1°E | 25 | 38902 | 52 | 92 |
| Liangshui | 47.1°N,128.2°E | 10 | 34021 | 44 | 82 |

**Table S3** Description of metadata used in the study. Some variables are not available for all samples (1453) and the availabilities (number of samples with available metadata) are noted in the table.

| Type | Abbreviation | Descripts | units | Availability |
|---|---|---|---|---|
| Host | dbh1 | the diameter at breast height | cm | 1418 |
| | LA | plant leaf area | $cm^2$ | 1070 |
| | LDMC | leaf dry matter content | % | 823 |
| | LeafC | leaf carbon content | % | 1009 |
| | LeafC.N | leaf carbon and nitrogen ratio | / | 1015 |
| | LeafN | leaf nitrogen content | mg/g | 1245 |
| | LeafP | leaf phosphorus content | mg/g | 1245 |
| | SLA | specific leaf area | $cm^2/g$ | 1253 |
| | WD | wood density | $g/cm^3$ | 1127 |
| Neighborhood | abund.10 | the abundance of plants within 10 meters | / | 1453 |
| | density | plant density in the plot | No./ha. | 1453 |
| | relativeabundance | the relative abundance of host plant species in the plot | / | 1453 |
| | richness.10 | the number of plant species within 10 m | / | 1453 |
| | sesmntd10 | Standardized effect size of mean nearest taxon distances of plant community within 10 m | / | 1453 |
| | sesmpd10 | Standardized effect size of mean pairwise distances in plant community within 10 m | / | 1453 |
| | spdensity | the density of host plant species | No./ha. | 1453 |
| Spatial | gx | the spatial coordinates of host plants in the local plot | m | 1453 |
| | gy | the spatial coordinates of host plants in the local plot | m | 1453 |
| | INTPTLAT | the geographic coordinates of the plot | degree | 1453 |
| | INTPTLONG | the geographic coordinates of the plot | degree | 1453 |
| Edaphic | Soil.AP | soil available phosphorus content | mg/kg | 1453 |
| | SoilACu | soil available cuprum content | mg/kg | 611 |
| | SoilAFe | soil available cerrum content | mg/kg | 611 |
| | SoilAK | soil available potassium content | mg/kg | 1453 |
| | SoilAN | soil available nitrogen content | mg/kg | 1453 |
| | SoilAZn | soil available zinc content | mg/kg | 611 |
| | SoilMn | soil available manganese content | mg/kg | 307 |
| | SoilNH4 | soil ammonium content | mg/kg | 239 |
| | SoilNO3 | soil nitrate content | mg/kg | 239 |
| | SoilOrgan | soil organic matter content | g/kg | 1453 |
| | SoilpH | soil pH | / | 1453 |
| | SoilTK | soil total potassium content | g/kg | 700 |
| | SoilTN | soil total nitrogen content | g/kg | 1453 |
| | SoilTP | soil total phosphorus content | g/kg | 1453 |
| | SoilVW | soil volume weight | g/cm3 | 649 |
| | Soilwater | soil water content | % | 1453 |
| Topo-graphic | Aspect | the slope aspect, cosine transformed | / | 917 |
| | Convex | surface convexity | radians/m | 1453 |

| | | | | |
|---|---|---|---|---|
| | Elev | altitude | m | 1453 |
| | Slope | topographic slope | degree | 1453 |
| Climates | aiet0 | global aridity index | / | 1453 |
| | BIO1 | annul mean temperature | centigrade | 1453 |
| | BIO12 | annul precipitation | mm | 1453 |
| | BIO15 | Precipitation seasonality i.e. coefficient of variation | / | 1453 |
| | BIO3 | temperature isothermality | / | 1453 |
| | et0 | potential evapotranspiration | / | 1453 |
